# Wintering, rather than breeding, oceanic conditions may modulate declining survival in a long-distance migratory seabird

**DOI:** 10.1101/2023.11.10.566398

**Authors:** Katherine R. S. Snell, Inês Alexandre Machado dos Santos, Rob SA van Bemmelen, Børge Moe, Kasper Thorup

**Affiliations:** Center for Macroecology, Evolution and Climate, Globe Institute, University of Copenhagen, Universitetsparken 15, DK-2100 Copenhagen, Denmark; MPI-AB, Am Obstberg 1, Radolfzell, 78315, Germany; Centre for the Advanced Study of Collective Behaviour, University of Konstanz, Konstanz, 78464, Germany; Waardenburg Ecology, Culemborg, 4101 CK, the Netherlands and Wageningen Marine Research, IJmuiden, 1976 CP, the Netherlands; NINA, Trondheim, 7485, Norway; Copenhagen Bird Ringing Centre, Natural History Museum of Denmark, University of Copenhagen, Øster Voldgade 5-7, DK-1350 Copenhagen, Denmark

**Keywords:** mortality, carry-over effects, CMR modelling, *Stercorarius parasiticus*

## Abstract

Steep declines in Arctic skua populations in the southern extent of their breeding range have been reported during the last half of the 20^th^ century. We used 24 years of ringing and re-encounter data from the Faroe Islands, North Atlantic, to investigate if patterns in survival probabilities can be explained by large scale climatic events. Having first determined the migratory phenology and wintering regions, we estimated the effects on survival of the North Atlantic Oscillation (NAO) index during breeding and Oceanic Niño index (ONI) during the non-breeding period within a capture-mark-recapture framework. Temporal trends, and direct and time-lagged effects of the environment on annual survival were modelled. We found support for a substantial decrease in adult annual survival, from ca. 0.93 in 1985 to ca. 0.77 in 2008, and weak support for a decrease in young (1^st^ year) survival over the duration of the study period. Furthermore, models indicated increased young survival following an El Niño winter. We suggest this time-lagged effect reflects downstream propagation of environmental conditions, particularly food availability, or a potential carry-over effect of El Niño conditions positively impacting the performance of the parents in the subsequent breeding season, leading to improved young survival prospects. While adult mortality cannot be attributed to the oceanic climate oscillations tested here; the negative trend in survival may account for the substantial population declines observed over the last decades.

## 1. INTRODUCTION

Seabirds occupy the upper trophic level and are important in island and marine ecosystem processes, function and resilience (Paleczny et al. 2015). Their distribution and population dynamics are strongly connected with their feeding ecology and patterns of food availability which are dependent on climatic and oceanographic factors, such as water temperature and currents (Meier et al. 2017). Therefore, changes in any such environmental variables can substantially influence seabird survival and productivity (Irons et al. 2008, Waugh et al. 2015, Champagnon et al. 2018). Steep declines in seabird numbers have been reported during the last half of the 20th century. Palenczny et al. (2015) assessed the population trend of the world’s monitored seabirds from a global dataset representing ca. 19% of the global seabird population and found an overall decline of 69.7% in between 1950-2010: a pattern reflected in the population trends of seabirds in the North Atlantic (Irons et al., 2008). Moreover, population declines are more apparent in migratory species (Kubelka et al. 2021).

Given that a broad range of factors may affect demographic parameters, identifying the factors responsible for population dynamics is often difficult. Unravelling effects of different factors is especially challenging for migratory species, as they depend on multiple, widely separated areas that may contrast in their (rate of change in) conditions. As migratory species spend the majority of their annual cycle on migration or overwintering, the habitat conditions experienced during these periods are likely to influence survival of all age classes post-fledging. Likewise, conditions on breeding grounds influence reproductive success, with decreases in productivity in years where conditions are poorer (Fife et al. 2018). In addition to these within-season effects, conditions experienced in one season can affect the condition and performance of individuals in later seasons (Harrison et al. 2011). The effects of climate on species survival have received increased interest during the last few decades due to accelerated climate change (Jóhannesson et al. 1995, Ballerini et al. 2015, Salas et al. 2017). As shown in several studies, climate indices can influence survival and breeding success of both migrant and non-migrant birds (England 2000, Irons et al. 2008, Boano et al. 2010, Genovart et al. 2013, Waugh et al. 2015). For example, blue-footed booby (*Sula nebouxii*) in the Gálapagos suffered increased mortality during an El Niño year, when sea surface temperature (SST) was elevated; similarly, chick growth rates of dark-rumped petrel (*Pterodroma phaeopygia*) were retarded during the 1983 nesting season, one of the strongest El Niño events since records began (England 2000). Formal analysis of long-term data revealed a complex interaction of stochastic oceanic processes on chick quality of sooty shearwaters (*Puffinus griseus*), including a time-lagged effect of El Niño oscillations (Humphries & Möller 2017). In adult blue petrels (*Halobaeria caerula*), Barbraud and Weimerskirch (2003) found that increased mortality occurred when the El Niño Southern Oscillation phase was associated with warmer water temperatures. The influence of climate indices on seabird survival rates are likely explained by the important impact of stochastic ocean scale environmental processes on ecosystem cascades affecting nutrients, temperature and primary productivity (Polis et al. 1997). Formally linking survival and reproductive rates to climate indices, such as the El Niño Southern Oscillation (ENSO) and North Atlantic Oscillation (NAO), can be a first step towards understanding the ultimate or proximate drivers of population dynamics (Descamps et al. 2010).

The influence of the ENSO in the Pacific Ocean triggers a cascade of events globally across equatorial regions and the Southern Ocean, with delayed propagation of conditions across the Atlantic, albeit less robust (Kim et al. 2023). During El Niño periods, warm Pacific waters are associated with increased atmospheric stability and suppressed tropical cyclone genesis over the Atlantic; while during La Niña years, cold equatorial Pacific SSTs are associated with decreased stability and increased tropical cyclones in the Atlantic between *ca.* 0-30°N (Kim et al. 2023). In the year following an El Niño event, warm water in the Atlantic may contribute to increased hurricane formation from Cape Verde towards the Americas (Kim et al. 2023). Overall higher wind speeds and lower SSTs around Atlantic upwellings are associated with significant increases in primary productivity (chlorophyll *a*) (Oviatt et al. 2015), which likely propagates into higher trophic levels. In contrast, periods of above average SSTs leads to reduced egg and fish abundance (Franco et al. 2020). The NAO index is simply the pressure differential between Iceland and the Azores. In the North Atlantic region the positive phase is related to warmer/wetter ambient conditions and more severe storms (Hurrell et al. 2001, Báez et al. 2021). Milder weather conditions lead to an advanced reproductive phenology (Báez et al. 2021), while increased precipitation and extreme weather events may have a direct impact on higher mortality (Stenseth et al. 2002, Frederiksen et al. 2008), and an increased energetic cost of migration (Nourani et al. 2023). Negative NAO phases are linked to relatively warmer water, cooler air temperature but dry/stable summers. NAO influences zooplankton abundance; and reflects fish both abundance and body condition, with fish larvae particularly sensitive to variation in water conditions (Stenseth et al. 2002, Drinkwater et al. 2003). In the waters east of the Faroes, NAO has a significant positive relationship with lower trophic levels: phytoplankton and zooplankton (Drinkwater et al. 2003). Around the Faroes, overall krill abundance was attributed to temperature and chlorophyll *a* (Silva et al. 2013); and the positive phase of the NAO is favourable for cod stocks, however the opposite is found in the North Sea, although the effect size is small (Drinkwater et al. 2003, Overland et al. 2010). Due to intense fishing practices there are difficulties in attributing prey demographics to oceanic processes in many regions (Oviatt et al. 2015). Cory’s shearwaters (*Calonectris borealis*) increased their foraging effort in negative phase NAO years, which was attributed to lower marine productivity in the eastern mid-North Atlantic (Paiva et al. 2017). In Scotland, NAO phase is a predictor of breeding phenology in black-legged kittiwakes (*Rissa tridactyla*) and common guillemots (*Uria aalge*), and breeding success of northern fulmars (*Fulmarus glacialis*) and subsequent population trajectories (Thompson & Ollason 2001, Frederiksen et al. 2004). While NAO clearly can be an important factor in seabird ecology, there are difficulties in making specific predictions for a given population.

Indices of large-scale ocean processes reflect multiple variations in the marine environment, including water and air temperatures, winds, salinity, run-off, and precipitation, and capture more of the environmental stochasticity than individual metrics. In general the influence of the oceanic phase is likely to be cumulative and dynamic and lags can be expected for some effects (Stenseth et al. 2002). While the underlying bottom-up mechanisms are complex and geographically explicit, they provide a valuable tool to elicit variation in biological processes.

Like many populations of vulnerable seabird species in Northwest Europe, (Dias et al. 2019), Arctic skua (*Stercorarius parasiticus*) populations are declining rapidly (Perkins et al. 2018). The Arctic skua is a migratory seabird with a circumpolar breeding range in high temperate to high Arctic zones. It is long-lived, with immature birds recruiting to the breeding population from an age of 4 years (O’Donald 1983, Furness 1987). They are kleptoparasitic and, thus, highly dependent on other seabirds (Furness 1987), most of which have also experienced sharp population declines (BirdLife International 2016). Although the Arctic skua is listed as ‘least concern’ globally, with a world population estimated at 4-600 000, it is considered ‘endangered’ within the European Union (Burfield & van Bommel 2004). In the United Kingdom the species has declined by *ca.* 66% during the period 2015-2021 (Burnell et al. 2023), and in the Faroe Islands the population declined from 1200-1500 pairs in 1981 (Bengtson & Bloch 2003) to *ca.* 207 pairs in 2017 (Santos 2018). The proposed drivers of these declines are geographically limited but, in the United Kingdom and Norway, associations were found with food shortages and predation of eggs and young (Perkins et al. 2018, van Bemmelen et al. 2021).

The decline of European Arctic skua populations has hitherto been mainly attributed to factors during the breeding periods on reproductive success, and adult survival was considered to be sufficiently high to maintain a stable population. However, a full understanding of the population declines requires quantification of potential effects from both the breeding and non-breeding periods, on both adult and immature mortality (Flack et al. 2022). We hypothesise that for long-lived, long-distance migratory seabirds non-breeding environmental conditions, rather than breeding conditions, influence survival rates and thereby population dynamics. Population rates for these life history traits are expected to be most sensitive to adult survival (Morris et al. 2011), as such are generally buffered against variations in environmental conditions experienced (Hilde et al. 2020). However, extreme weather events influence survival (Frederiksen et al. 2008) and are likely to be encountered more often during the non-breeding period, when conditions are less predictable (Lisovski et al. 2017). Here, we investigated annual survival of adult and young Arctic skuas breeding in the Faroe Islands. Within a formal capture-mark-recapture framework we explored annual survival, trends in annual survival and the influence of oceanic scale climatic events, precisely the El Niño Southern Oscillation (ENSO) and North Atlantic Oscillation (NAO), using a ringing dataset spanning over two decades. We predict that non-breeding conditions, the largest component of the annual cycle (van Bemmelen et al. 2024), will influence annual survival of both young and adults, with El Niño years having an overall negative effect due to increased water temperatures leading to reduced marine productivity. Furthermore, in long-lived species, we expect adult survival to be largely buffered against environmental variability; as such, mortality will firstly be greater in young than in adults, and secondly, instead of only affecting survival, poor feeding conditions may be expected to carry over to the breeding performance due to the adults’ poorer condition upon arrival and a subsequent increased mortality of offspring.

## 2. MATERIALS & METHODS

### 2.1 Study Species and sites

The Faroe Islands (61°53’N, 6°58’W) are located between the UK and Iceland, in the middle of the North Atlantic current which creates stable temperatures and a maritime climate (Fosaa et al. 2004). The Faroese archipelago consists of 18 islands and is an important breeding site for seabirds (Bakken et al. 2006). Ringing of Arctic skuas has been conducted since 1924 across the archipelago but was not systematic (Hammer et al. 2014).

As part of a tracking programme covering the Atlantic AMAs (Arctic Marine Areas) (Kuletz et al. 2017), tracking data was obtained from 20 birds in the Fugloy colony from 2016-7 (Fig. 1). Full details of the analysis and interpretation of light level data are detailed in van Bemmelen et al. (2024). The Faroese Arctic skua population winter on the Patagonian Shelf, Discovery Seamounts, coastally from the Iberian plain to Cape Verde Islands, the Gulf of Guinea, and off the Namibian coast and the Cape of Good Hope (Fig.1). Spatio-temporal data was used to select appropriate environmental indices. Mean arrival and departure dates were 27 April and 8 August for breeding and 9 September and 21 March for over-wintering sites, respectively.

**Fig. 1.**
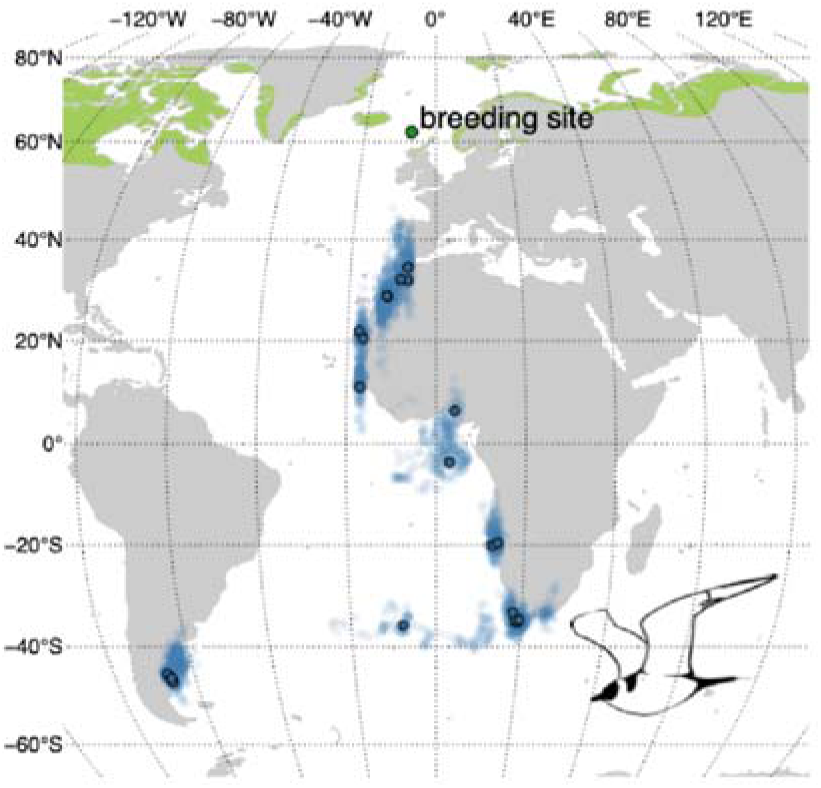
Non-breeding areas of Arctic skuas breeding in Fugloy, Faroe Islands, based on positional data from 20 GLS tracked birds 2016-7 (van Bemmelen et al., 2024): blue transparent circles are daily wintering GLS positions and blue circles with black outlines indicate the centroid of each bird. Birds largely over-winter on the Patagonian Shelf, Discovery Seamounts, coastally from the Iberian Peninsula to the Cape Verde Islands, the Gulf of Guinea, the Angola Current and from the Benguela Current westwards to Tristan da Cunha. The breeding range is given in green and the Faroe Islands study site is indicated by the dark green circle. Breeding range data from BirdLife International and Handbook of the Birds of the World (2016); a Robinson projection was used to create the map.

### 2.2 Environmental Variables

Standardised large-scale oceanic indices were selected appropriate for a spatio-temporal schedule of the long-distance migrant. The Oceanic Niño index (ONI) quantifies the phase and strength of El Niño Southern Oscillation (ENSO) events, where negative values correspond to La Niña years and positive values to El Niño years (NOAA 2018a). The North Atlantic Oscillation index (NAO) is the air pressure differential across the North Atlantic and represents the phases of the North Atlantic Oscillation (NOAA 2018b). Both ONI and NAO data are presented as aggregated 3-month overlapping temporal periods with values ranging from -1.64 to 2.24 and -1.4 to 1.8, respectively (Fig. 2). Wintering and breeding periods were derived from the GLS data. We calculated mean ONI during the stationary non-breeding season (September to March) and mean NAO during the breeding season (May to July; Fig. 2). Values were used as covariates in the survival model both as a direct effect (same year) and as a time-lagged effect from the previous year.

**Fig. 2.**
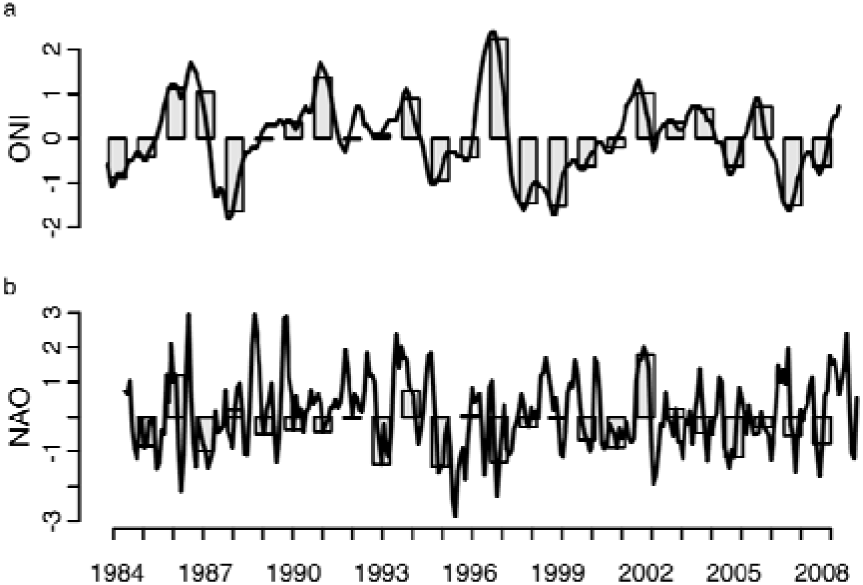
a. The Oceanic Niño index (ONI) and b. North Atlantic Oscillation index (NAO). Lines represents 3-month rolling mean per calendar month. Bars shows the mean aggregate for period of interest: a. non-breeding period (September to March) and b. breeding period (May to July), of which values were used as covariates in the survival models.

### 2.3 Survival Analysis

We used a subset (1985 to 2008) of the ringing and recovery data which had been digitised and had the greatest continuity of data (prior to 1985 gaps of >2 years in ringing effort occurred). Where discrepancies made the record unreliable, we excluded these individuals from our dataset (n=2). The dataset totalled 1060 individuals, of which 530 were ringed as chicks and 530 ringed as adults; 85 individuals were subsequently re-encountered (alive and/or dead) during the 24 years (suppl. info. Fig. S1). Live re-encounters consisted of ring reading during breeding season fieldwork; dead recoveries included both individuals recovered from the study colony and rings reported to the ringing scheme. The reporting probability structure is similar to the ringing effort for the period included in this study. A year was defined as starting on the 1^st^ May and ending on the 30^th^ April of the following calendar year, to coincide with the birds’ breeding cycle. For young birds the initial year is the period from ringing (typically from 10 days after hatching) until the following breeding season – and therefore includes late chick stage, fledging, and dependant period.

Annual survival was estimated from a formal capture-mark-recapture framework with the Mark software, version 10.1 (White & Burnham 1999). Firstly, we use the Burnham model for joint live and dead encounters (Burnham 1993). This method calculates parameter estimates of apparent survival (S), live re-encounter probability (p), dead recovery probability (r) and fidelity (F). An encounter history was created for each individual in the classic 2 age-class structure: for young (1^st^ year) and for adults and included both live and dead encounters. We modelled fidelity as constant due to known high philopatry and no reported post-natal dispersal between the Faroe Islands and the nearest colonies in the UK (Hammer et al. 2014): F was found to be high, F=0.94 +/-0.065 SE. As Arctic skuas generally delay breeding until their 4^th^ season (O’Donald 1983, Furness 1987), re-encounter (p) and recovery probability (r) were modelled as a function of 3 age-classes: young, immature (Imm) and adults. During the immature years (age 1 to 3 inclusive), birds are thought to be entirely marine, not returning to land once they leave the colony (Furness 1987); as such p(Young), p(Imm) and r(Imm) were fixed to 0. As ringed young birds could be recovered dead in the colony after the breeding season had finished r(Young) was estimable (there were three occurrences in the data where birds ringed as chicks were recovered dead within the first year). Survival probability (S) was modelled as a function of 2 age-class (Young and Adults) with the assumption that once birds are full-grown immatures, after 1 year, they will have the same mortality as breeding adults. As the joint model includes more parameters than a live-only re-encounter model we also repeated the analysis by modelling apparent survival with a Cormack-Jolly-Seber (CJS) model (Lebreton et al. 1992). This method calculates parameter estimates of apparent survival (Φ) and live re-encounter probability (p). The encounter history for each individual was created in the same 2 age-class structure: for young (1^st^ year) and for adults using only live encounter data. The structure of the model followed the Burnham model, and p(Young) and p(Imm) were fixed to 0.

We derived a basic model (i.e., a single survival probability over all years) and modelled year-dependent survival for each age-class. We then compared the following *a priori* candidate models: linear temporal trends of survival rates for both age-classes combined and for each age-class independently, and the effects of the two environmental covariates (detailed above) on survival probabilities. We estimated a dispersion coefficient with the bootstrap goodness-of-fit function in MARK to the general model and some over-dispersion was indicated. Residuals were visually inspected and minor departures from goodness-of-fit did not indicate a structural issue. Therefore, we calculated the variance inflation factor (ĉ) via the median c-hat procedure in MARK, and then used this to adjust AICc values (Akaike 1973). We tested the overall CJS model encounter histories using the overall_CJS() function in the R package R2ucare (Gimenez et al. 2018; a formal test for the Burnham model is not available). This tests the basic assumptions of that model and indicated the CJS models were appropriate for the data and any overdispersion in the data was not significant (χ^2^ = 16.181, p = 0.973, df = 29). No formal statistical method for testing goodness-of-fit (GOF) for models with linear covariates (here, trends over time or environmental covariates) is available. Model selection was based on the ranking of Akaike’s information criteria corrected for small sample size adjusted for median-ĉ (AICc). Survival estimates ± SE are given as back transformed from the logit link function Beta parameter estimates. Linear trends are visualised with 95% confidence intervals.

## 3. RESULTS

The basic joint Burnham model (model B, Table 1) estimated the overall survival of young as 0.43±0.14 and adults 0.92±0.07. Year dependant annual survival was estimable in 10 of the 24 years for adults, ranging from 0.16±0.15 to 0.87±0.51, and in 8 years for young, ranging from 0.13±0.12 to 0.66±0.67 (the time-dependant model is not included in Table 1 due to the high number of non-estimable parameters).

**Table 1.** Annual survival model results for Arctic Skuas, relative to the highest ranked model [C1] and grouped into temporal trend models [T1-3] and environmental variable models [E1-6]. The basic model is given by [B]. Akaike’s information criterion values adjusted for median-ĉ; (AICc), differences in AICc (ΔAICc), within-group ΔAICc, weights, model likelihood, number of estimable parameters and deviance are given. 2Age represents 2 age-classes. We present the best supported environmental indices: ONI(t-1) is the time lagged (-1 year) Oceanic Niño index, NAO is the North Atlantic Oscillation index. The three equally supported best models are highlighted by bold ΔAICc. Live re-encounter probability (p), dead recovery probability (r) and fidelity (F) remain constant across models: p(3Age_Young=Imm=0_), r(3Age_Imm=0_), F(.).

| | Model | AICc | ΔAICc | Within-group ΔAICc | AICc Weights | Model Likelihood | no. Parameters | Deviance |
| --- | --- | --- | --- | --- | --- | --- | --- | --- |
| C1 | 2Age + Trend<sub>Adult</sub> + ONI(t-1)<sub>Young</sub> | 2383.7 | **0.00** | 0.00 | 0.245 | 1.00 | 8 | 1772.72 |
| B | 2Age | 2387.2 | 3.56 | 3.56 | 0.041 | 0.17 | 6 | 1780.34 |
|  | **Temporal Trend Models** |  |  |  |  |  |  |  |
| T1 | 2Age + Trend<sub>Adult</sub> | 2384.4 | **0.70** | 0.00 | 0.173 | 0.71 | 7 | 1775.45 |
| T2 | 2Age + Trend<sub>2Age</sub> | 2385.1 | **1.38** | 0.68 | 0.123 | 0.50 | 7 | 1776.13 |
| B | 2Age | 2387.2 | 3.56 | 2.86 | 0.041 | 0.17 | 6 | 1780.34 |
| T3 | 2Age + Trend<sub>Young</sub> | 2389.0 | 5.32 | 4.62 | 0.017 | 0.07 | 7 | 1780.07 |
|  | **Environmental Variable Models** |  |  |  |  |  |  |  |
| E1 | 2Age + ONI(t-1)<sub>Young</sub> | 2385.9 | 2.19 | 0.00 | 0.082 | 0.33 | 7 | 1776.94 |
| B | 2Age | 2387.2 | 3.56 | 1.37 | 0.041 | 0.17 | 6 | 1780.34 |
| E2 | 2Age + NAO<sub>Adult</sub> | 2388.3 | 4.58 | 2.39 | 0.025 | 0.10 | 7 | 1779.34 |
| E3 | 2Age + ONI(t-1)<sub>Adult</sub> | 2388.5 | 4.84 | 2.65 | 0.022 | 0.09 | 7 | 1779.59 |
| E4 | 2Age + NAO<sub>Young</sub> | 2389.3 | 5.59 | 3.40 | 0.015 | 0.06 | 7 | 1780.34 |
| E5 | 2Age + ONI(t-1) | 2393.8 | 10.1 | 7.91 | 0.001 | 0.01 | 6 | 1786.92 |
| E6 | 2Age + NAO | 2397.9 | 14.2 | 12.01 | 0.001 | 0.01 | 6 | 1791.02 |

Three models were well supported (ΔAICc < 2, Table 1; model C1, T1 & T2). The highest ranked model (model C1), indicated a combined negative trend for adult annual survival and the influence of a time-lagged effect of ONI (Oceanic Niño Index) on young survival (Table 1; suppl. info. Table S1 for parameter estimates); it should be noted that this model is nested. Evidence for a negative trend in adult annual survival is further supported by model T1 (ΔAICc = 2.86 from model B, the basic model (Table 1; suppl. info. Table S2 for parameter estimates)). We found some support for a decreasing trend for young birds also (Table 1; suppl. info. Table S3 for parameter estimates). Based on model T2, the adults’ survival showed a decline of 17% in 24 years, with 0.93±0.05 probability of survival in 1985 and 0.77±0.09 in 2008, whereas young survival decreased 57%, from 0.60±0.19 in 1985 to 0.26±0.14 on 2008 (Fig. 3a,b). Adult survival estimates from model C1 and T1 were comparable: within the error range of model T2 estimates.

**Fig. 3.**
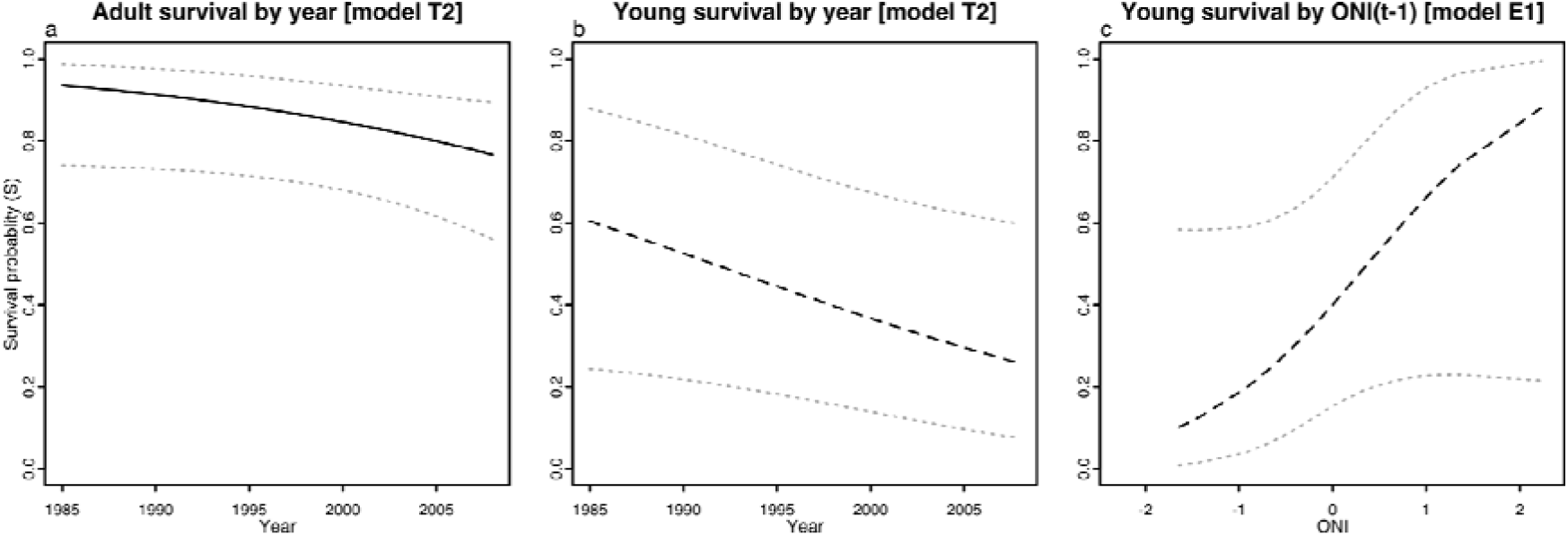
Best supported models describing variation in survival probabilities for (a) adult and (b-c) young. Plots represent the back transformed (logit link function) derived linear trend with (a-b) year (Table 1 [model T2]; suppl. info. Table S3) and (c) the time-lagged oceanic Niño index, ONI(t-1) (Table 1 [model E1]; suppl. info. Table S4), with 95% confidence intervals. Solid lines indicate strongly supported models, dashed lines are weakly supported (i.e. there is equal support for no trend: Table 1 [model T1]).

Model support indicated that young survival could be influenced in part by ONI from the preceding year. El Niño events were related to an increase in apparent survival while La Niña events a reduction (Fig. 3c & Table 1). When testing for the effect of time-lagged ONI on young in isolation (model E1; suppl. info. Table S4 for parameter estimates) the model was only weakly supported, suggesting a coupled interaction with a temporal trend. The model which included temporal trends for both age-classes and ONI carry-over effect on young survival included several non-estimable parameters and was not supported. No models including the NAO (North Atlantic Oscillation) during breeding season as a covariate were supported (Table 1; Group E).

The live-only CJS model corroborated results from the joint Burnham model, firstly with a similar pattern in apparent survival in the basic model. The highest ranked model was the same: a trend in adult survival and time-lagged ONI effect on young (suppl. info. Table S5 & S6). Model ranking was equivalent when using a ΔAIC of <2 to indicate equal support (suppl. info. Table S5). Because of the limited number of live young in the CJS model, we focus on results from the Burnham model.

## 4. DISCUSSION

Survival rates and productivity are important parameters in population dynamics and, in long-lived species, survival probability is the greatest contributing factor (Lebreton & Clobert 1991). The numerous ecological factors that are acting upon survival rates, and the tendency for survival to be buffered against environmental stochasticity in long-lived species, complicate determining the driving factors. However, survival rates of seabirds are known to be affected by climatic events, and increased variation in environmental conditions leads to increased variation in vital rates (Morris et al. 2011). In our study of Arctic skuas breeding on the Faroe Islands, a declining trend in survival of adult birds with a possible decline in survival of young birds (1^st^ year) coupled with a time-lagged/carry-over effect of ENSO best explained the variation in the data. These findings may in part explain the notable decline of the Arctic skua population in the Faroe Islands in recent times, and should be considered for other seabird species experiencing similar declines.

As K-selected species, many seabirds share the life history trait of high annual survival for adults (Szostek & Becker 2015). Overall high survival probabilities of this age-class are commensurate with reported values and are expected for Arctic skuas (O’Donald 1983, Phillips 2001, Davis et al. 2005, van Bemmelen et al. 2021). We found strong support for a negative trend over the 24-year study period. Because annual survival of adults is important for maintaining a stable population, a decrease of 17% in survival probability in this timeframe is certainly biologically significant to population persistence (van Bemmelen et al. 2021). Due to their demographic strategy, such a significant decline in adult survival is both unusual and alarming. In general, in long-lived vertebrates, the rate of population growth is most sensitive to adult survival. The number of Arctic skuas breeding in the Faroe Islands has decreased substantially during our study period (Santos 2018) and the negative trend in adult survival observed on our study might explain this decline. While the cause of increased mortality in this population has yet to be determined, the potential impact on the viability of this population is concerning. In adults, we found no support for the influence of large-scale oceanic indices on their survival, either during the breeding season or non-breeding period. This may be due to the masking of any effect by the strong downward trend, or be due to an environmental variable trending over the duration of the study period (such as air/sea surface temperature) as it would be difficult to resolve drivers of the decline over collinearity (Grosbois et al. 2008), or limitations in the data. As such, exploration of this pattern demands further investigation, aided in part by our improved understanding of the spatial ecology of this population and species. The closest neighbouring population, in the UK, is also suffering heavy population declines (Jones et al. 2008, Perkins et al. 2018). Here, studies aimed at understanding these population declines have focussed mainly on explaining population dynamics by the variation in reproductive success (Perkins et al. 2018), but have left open the possibility that population declines were (also) driven by reduced survival rates due to factors outside the breeding period. Furthermore, Perkins et al. (2018) demonstrated that low breeding success affects the stability of the population. The breeding success of Arctic skuas in the Faroe Islands has received no formal analysis to date, but sharp declines of the population are expected during our study period if productivity declined here also, together with the decreases in adult survival observed in our study.

ENSO has been linked with survival in various seabird species (Boersma 1998, England 2000, Champagnon et al. 2018). These examples, however, only assessed chick survival from birth to fledging. In the Kiritimi Island Black noddy (*Anous minutos*) populations, no nestlings survived during an El Niño year, when heavy rains flooded the nests, and in the Galapagos archipelago reduced chick size and survival was reported in three species during an El Niño year (Jaksic & Fariña 2010). While we found no direct effect of ENSO on adult survival, we did find some support for an influence of ENSO conditions (ONI) from the previous year on first year survival (specifically in the period from ringing until the following breeding season). One explanation is a possible carry-over effect observed on young birds that might be a consequence of the quality of their parents’ non-breeding environment. Alternatively, a time-lagged response may be attributed to downstream ecological cascades. A time-lagged effect is commonly reported in relation to life cycles of prey species as the influence of the environment is propagated up the food chain (Stenseth et al. 2002). As such, the effect of conditions during the previous year may indicate that survival of young skuas is governed by food availability. Szostek and Becker (2015) found such a correlation between primary productivity during winter with recruitment in the following breeding season in common terns (*Sterna hirundo*). Primary productivity can be used as a proxy for food availability (Lindeman 1942) and this suggests that food availability during the non-breeding period is limiting survival and recruitment success in common terns. As ENSO has effects on primary productivity, ENSO would be expected to correlate with species vital rates, as indicated in our study. Potentially, in Arctic skuas, as in the case of common terns, the quality of the environment during the non-breeding period for adult birds, expressed by food availability, will be reflected on the quality of their young and their ability to complete their first year. There is likely to be a disparity between the impact of south Atlantic conditions on locally breeding populations and migratory birds from thousands of kilometres away. Alternatively, the time-lagged influence of ONI may result from the direct interaction of a delayed effect of environmental processes in the Atlantic from ENSO phenomena originating in the Pacific; specifically, increased wind associated with El Niño events in the previous year around important upwellings where skuas are known to winter (Iberian and Benguela currents), and increased primary production, may lead to more profitable foraging (Oviatt et al. 2015, Kim et al. 2023). Although our model support for an effect of the wintering environment was not strong, we did find some support for a downward trend in survival probability in young over time (the model including constant survival probability was equally supported with the temporal trend). The coupled interaction of the stochastic ENSO events masking the trend over time may explain the limited model support. However, the magnitude of the decline is substantial (57%), as such the potential biological significance of lowered recruitment cannot be ignored over and above a potential interaction with ONI and/or the temporal occurrence of ringing data (see below). In terms of quantifying higher-level processes, we acknowledge that while the trend is indicated, the magnitude of the effect is uncertain (i.e., large confidence intervals around the estimate). For such vulnerable populations, employing alternative analysis, such as integrated population models, may improve the robustness of estimates (Schaub & Abadi 2011).

We found no evidence of the effects of NAO on survival of either adults or young. This suggests the lack of a strong influence of oceanic conditions on the survival of both young and adults during the breeding season. One explanation is that all skuas, regardless of age spend the majority of their annual cycle in the wintering grounds rather than the breeding region in the north Atlantic (Hurrell et al. 2001, Stenseth et al. 2003), and so have more limited exposure to the effects of NAO. Furthermore, the summer season is characterised by clement weather and abundant resources which is likely to buffer the influence of oceanic processes; in addition, the influence of NAO effects are regionally specific and in our area the magnitude and direction of the effect on fish abundance, and hence prey availability is variable (Drinkwater et al. 2003, Overland et al. 2010, Báez et al. 2021). It is likely that the environmental conditions during the breeding season (and the influence of NAO) have greater effects on breeding success than on survival (Thompson & Ollason 2001, Frederiksen et al. 2004, Frederiksen et al. 2007).

While we made attempts to use the best available dataset to cover the range of stochastic environmental conditions (in particular intense El Niño and La Niña events) and this modelling approach takes into account a lack of data in the estimates, the structure of the ringing and re-encounter data precluded a fully time-dependent model (year by year probability estimates) for all years of the study period. This limits our ability to detect an effect of the linear covariates and influences how the results are interpreted. We conservatively state that the absence of an effect of the environment, consistent with the demographic buffering hypothesis (Hilde et al. 2020) cannot be ruled out. While we found a signal of an effect of time-lagged ONI on survival of young birds, AIC support was not definitive; furthermore, this alone does not indicate the environmental or ecological mechanisms potentially involved in driving this relationship.

Not all variation in survival could be attributed to the variables tested in this study. This may be explained by the complex interactions and regionally specific effects of stochastic oceanic events, or the inability of these fundamental models to fully capture nuanced ecological processes (including variation in other sources of mortality such as predation, hunting etc.). A better understanding of direct, carry-over, time-lagged effects of the climatic event or time-lagged trophic propagation is needed to describe the mechanistic processes at play. An exciting knowledge gap, yet to be explored due to the current lack of technology, is an understanding of the movements of young and immature birds in comparison to non-breeding movements of adults. Furthermore, population changes can be investigated with an approach from breeding ecology, which is outwith the scope of this survival study. Future studies should include the analysis of the contribution of other demographic parameters as a function of climate variables, to also help understand how climate influences survival from hatching until the chicks fledge.

## 5. CONCLUSIONS

From a conservation perspective, studying population dynamics and understanding the underlying process is important for targeting effective conservation measures. The declining trend on annual survival of adult birds observed in our study is an important component in understanding population declines in the Faroe Islands. Additionally, the time-lagged effect of ENSO on survival of young birds can inform predictions of population trends, especially those due to climate change effects and the prediction of an increase of ENSO events, despite our incomplete understanding of the complex interactions of ENSO environmental and ecological processes on survival rates. We highlight the importance of continuing ringing efforts and longitudinal monitoring programmes, particularly in light of acute conservation threats such as hunting (BirdLife International 2017) and highly pathogenic strains of avian influenza (Banyard et al. 2022).

## Supporting information

suppl. mat.

## ACKNOWLEDGEMENTS

Ringing data is courtesy of the Danish Bird Ringing Centre at the Natural History Museum of Denmark, University of Copenhagen. We are indebted to Willy Mardel for his long-term ringing efforts and his help resolving any data uncertainties in archival records. We gratefully acknowledge the support of Faroes Museum of Natural History and Leivu Janus Hansen, Sjúrður Hammer, Jón Aldará and Anthony Wetherhill for their substantial involvement in the GLS tracking project fieldwork. We kindly acknowledge the landowners on Fugloy for permission to work on their property. Catching and deploying geolocators was approved in the Faroe Islands by the Copenhagen Bird Ringing Centre and The National Museum of the Faroe Islands. A pre-print of this work is available at https:// https://www.biorxiv.org/content/10.1101/2023.11.10.566398v1. We kindly thank the anonymous reviewers for their comments which greatly improved the manuscript.

