## Supplementary material for "Wintering, rather than breeding, oceanic conditions may modulate declining survival in a long-distance migratory seabird": suppl. mat.

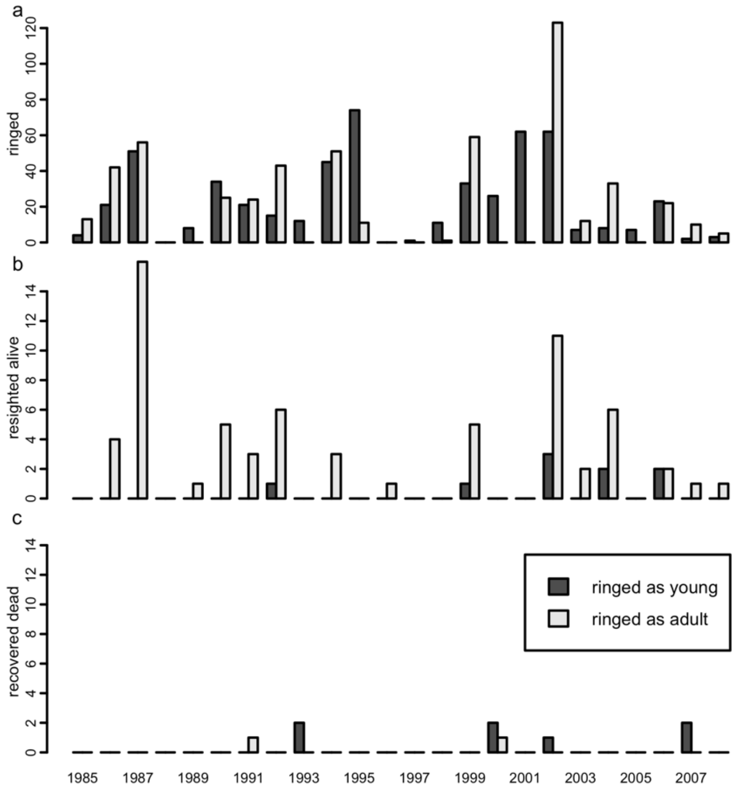


**Fig. S1**. Ringing and re-encounters from 1985 until 2008, by age-class ringed. a. Total number of ringed individuals; young birds are ringed as flightless chicks and adults are typically breeding birds caught on the nest. b. Total number of live re-encounters from 1985 until 2008. c. Total number of individuals recovered dead from 1985 until 2008.

**Appendix 1. Modelling process workflow**.

1. **Age Classes.** Survival probability was modelled as an annual estimate for two age classes (Young and Adults) with the assumption that once birds are full-grown immatures, after 1 year, they will have the same mortality as breeding adults.
2. **Basic Model.** A single survival probability generated over all years for Young and Adult
3. **GOF, Transience and Trap Dependence (CJS).** The Basic CJS model encounter histories was tested using UCARE.
4. **Linear covariates (Temporal Trends & Environmental Indices).** We included linear temporal trends of survival rates for both age-classes combined and for each age-class independently; and the effects of the two environmental covariates directly or as a time-lagged effect from the previous year (ONI, ONI(t-1), NAO, NAO(t-1)).

**Table S1.** Parameter estimates for model C1, which combined temporal trends and environmental covariates: S(2Age + $Trend_{Adult}$ + $ONI(t-1)_{Young}$), p($3Age_{Young=Imm=0}$), r($3Age_{Imm=0}$), F(.). This was one of the top three models which were equally supported (Table 1).

| Index | Label | Estimate | SE | LCI | UCI | |
|---|---|---|---|---|---|---|
| 1 | Intercept | 2.952853 | 1.024757 | 0.94433 | 4.961376 | |
| 2 | Age | -3.30956 | 1.170375 | -5.6035 | -1.01563 | |
| 3 | Adult trend | -0.07685 | 0.046505 | -0.168 | 0.014301 | |
| 4 | ONI (t-1) young | 0.975774 | 0.673861 | -0.34499 | 2.296542 | |
| 5 | Recapture young | -5.28786 | 1864.865 | -3660.42 | 3649.848 | Fixed |
| 6 | Recapture immatures | -0.24205 | 403.5296 | -791.16 | 790.6759 | Fixed |
| 7 | Recapture adults | -3.45144 | 0.186772 | -3.81751 | -3.08537 | |
| 8 | Recovery young | -4.60501 | 0.630517 | -5.84082 | -3.3692 | |
| 9 | Recovery immatures | -2.30189 | 979.2815 | -1921.69 | 1917.09 | Fixed |
| 10 | Recovery adults | -4.55623 | 0.423411 | -5.38612 | -3.72635 | |
| 11 | Fidelity | 2.786192 | 1.19165 | 0.450557 | 5.121826 | |

**Table S2.** Parameter estimates for model T1: S(2Age + $Trend_{Adult}$), p($3Age_{Young=Imm=0}$), r($3Age_{Imm=0}$), F(.). This was one of the top three models which were equally supported (Table 1).

| Index | Label | Estimate | SE | LCI | UCI | |
|---|---|---|---|---|---|---|
| 1 | Intercept | 3.0159022 | 1.053873 | 0.950311 | 5.081493 | |
| 2 | Age | -3.195031 | 1.157379 | -5.46349 | -0.92657 | |
| 3 | Adult trend | -0.082368 | 0.048113 | -0.17667 | 0.011933 | |
| 4 | Recapture young | -0.144142 | 0 | -0.14414 | -0.14414 | Fixed |
| 5 | Recapture immatures | -0.144363 | 0 | -0.14436 | -0.14436 | Fixed |
| 6 | Recapture adults | -3.455888 | 0.187155 | -3.82271 | -3.08906 | |
| 7 | Recovery young | -4.554322 | 0.641781 | -5.81221 | -3.29643 | |
| 8 | Recovery immatures | -0.144353 | 0 | -0.14435 | -0.14435 | Fixed |
| 9 | Recovery adults | -4.565737 | 0.42246 | -5.39376 | -3.73772 | |
| 10 | Fidelity | 2.8429757 | 1.250611 | 0.391778 | 5.294173 | |

**Table S3.** Parameter estimates for model T2: S(2Age + $Trend_{2Age}$), p($3Age_{Young=Imm=0}$), r($3Age_{Imm=0}$), F(.). This was one of the top three models which were equally supported (Table 1). Back transformed estimates were used to visualise trends in Fig. 3a,b.

| Index | Label | Estimate | SE | LCI | UCI | |
|---|---|---|---|---|---|---|
| 1 | Intercept | 2.737658 | 0.859846 | 1.05236 | 4.422956 | |
| 2 | Adult trend | -2.2485 | 0.821269 | -3.85819 | -0.63881 | |
| 3 | Young trend | -0.06448 | 0.036067 | -0.13517 | 0.006209 | |
| 4 | Recapture young | -0.02347 | 3E-07 | -0.02347 | -0.02347 | Fixed |
| 5 | Recapture immatures | -0.02347 | 4.2E-06 | -0.02348 | -0.02346 | Fixed |
| 6 | Recapture adults | -3.45447 | 0.18673 | -3.82046 | -3.08848 | |
| 7 | Recovery young | -4.59911 | 0.639979 | -5.85346 | -3.34475 | |
| 8 | Recovery immatures | 3.936313 | 0 | 3.936313 | 3.936313 | Fixed |
| 9 | Recovery adults | -4.54837 | 0.425205 | -5.38177 | -3.71497 | |
| 10 | Fidelity | 2.891914 | 1.372113 | 0.202572 | 5.581255 | |

**Table S4.** Parameter estimates for model E1 S(2Age + $ONI(t-1)_{Young}$), p($3Age_{Young=Imm=0}$), r($3Age_{Imm=0}$), F(.). Back transformed estimates were used to visualise trends in Fig. 3c.

| Index | Label | Estimate | SE | LCI | UCI | |
|---|---|---|---|---|---|---|
| 1 | Intercept | 2.4230526 | 1.0097595 | 0.4439239 | 4.4021813 | |
| 2 | Age | -2.8333182 | 1.1261593 | -5.0405905 | -0.6260458 | |
| 3 | ONI (t-1) young | 1.0755829 | 0.6801406 | -0.2574928 | 2.4086586 | |
| 4 | Recapture young | -0.1045219 | 0 | -0.1045219 | -0.1045219 | Fixed |
| 5 | Recapture immatures | 1.3047824 | 0 | 1.3047824 | 1.3047824 | Fixed |
| 6 | Recapture adults | -3.4774494 | 0.1850274 | -3.840103 | -3.1147958 | |
| 7 | Recovery young | -4.6200625 | 0.6260448 | -5.8471103 | -3.3930146 | |
| 8 | Recovery immatures | 1.7162662 | 0 | 1.7162662 | 1.7162662 | Fixed |
| 9 | Recovery adults | -4.2216151 | 0.6151004 | -5.4272118 | -3.0160184 | |
| 10 | Fidelity | 2.2758548 | 0.93235 | 0.4484488 | 4.1032609 | |

**Table S5.** Annual survival model results for Arctic Skuas comparing the joint live and dead data, Burnham model and apparent survival results for live re-encounter only data, CJS model for the top five supported models. All models are ranked in the same order and trends are in the same direction. We report differences in Akaike's information criterion values adjusted for median-$\hat{c}$; (AICc), and number of estimable parameters to allow direct comparison of modelling approaches. Model structures include temporal trends and environmental parameterization variables: 2Age represents 2 age-classes; ONI(t-1) is the time lagged (-1 year) Oceanic Niño index. Live capture probability (p) for Burnham and CJS models, and dead recovery probability (r) and fidelity (F) for Burnham models remain constant: p(3Age$_{Young=Imm=0}$), r(3Age$_{Imm=0}$), F(.).

§ Basic model

| | | **Burnham model – Live and Dead data** | | | **CJS model – Live only data** | | |
|---|---|---|---|---|---|---|---|
| | **Model** | **ΔAICc** | **Model Likelihood** | **no. Parameters** | **ΔAICc** | **Model Likelihood** | **no. Parameters** |
| 1 | 2Age + Trend$_{Adult}$ + ONI(t-1)$_{Young}$ | 0.00 | 1.00 | 8 | 0.00 | 1.00 | 5 |
| 2 | 2Age + Trend$_{Adult}$ | 0.70 | 0.71 | 7 | 2.00 | 0.37 | 4 |
| 3 | 2Age + Trend$_{2Age}$ | 1.38 | 0.50 | 7 | 3.98 | 0.14 | 5 |
| 4 | 2Age + ONI(t-1)$_{Young}$ | 2.19 | 0.33 | 7 | 2.36 | 0.30 | 4 |
| 5 | 2Age § | 3.56 | 0.17 | 6 | 5.18 | 0.08 | 3 |

**Table S6.** Parameter estimates for the best supported model using live only data (CJS): $\Phi$(2Age + Trend$_{Adult}$ + ONI(t-1)$_{Young}$), p(3Age$_{Young=Imm=0}$).

| **Index** | **Label** | **Estimate** | **SE** | **LCI** | **UCI** | |
|---|---|---|---|---|---|---|
| 1 | Intercept | 2.37768 | 0.45355 | 1.48872 | 3.26664 | |
| 2 | Age | -3.85439 | 0.8845 | -5.58805 | -2.12072 | |
| 3 | ONI (t-1) young | 1.489752 | 0.91806 | -0.309651 | 3.289156 | |
| 4 | Adult trend | -0.060072 | 0.02981 | -0.118513 | -0.00163 | |
| 5 | Recapture young | 0.00000 | 0.00000 | 0.00000 | 0.00000 | Fixed |
| 6 | Recapture immatures | 0.00000 | 0.00000 | 0.00000 | 0.00000 | Fixed |
| 7 | Recapture adults | -3.450204 | 0.183774 | -3.810402 | -3.090007 | |
